# Enhanced K_Na_1.1 Channel Underlies Cortical Hyperexcitability and Seizure Susceptibility after Traumatic Brain Injury

**DOI:** 10.1101/2023.10.14.562359

**Authors:** Ru Liu, Lei Sun, Le Du, Lulu Yu, Peng Pei, Xi Guo, Meng Jia, Qun Wang, Jianping Wu

**Affiliations:** Department of Pharmacology, Hubei University of Medicine, Shiyan 442000, China; Beijing Tiantan Hospital, Capital Medical University, Beijing 100070, China; China National Clinical Research Center for Neurological Diseases, Beijing 100070, China; Advanced Innovation Center for Human Brain Protection, Capital Medical University, Beijing 100070, China; School of Chemistry, Chemical Engineering and Life Sciences, Wuhan University of Technology, Wuhan 430070, China

**Keywords:** Traumatic brain injury, K_Na_1.1 channel, Seizure susceptibility, Neuronal hyperexcitability, imbalanced E/I

## Abstract

Pathogenic variants of the sodium-activated potassium channel K_Na_1.1 have been reported in multiple epileptic disorders. However, whether and how K_Na_1.1 channel is involved in epileptogenesis after traumatic brain injury (TBI) remains unknown. Firstly, we used behavioral monitoring and EEG recording to examine physiological property, spontaneous seizure activity, and seizure susceptibility after TBI. We explored the changes of K_Na_1.1 channel following TBI, including changes of subcellular distribution and expression pattern. Meanwhile, we performed patch-clamp recording to detect the neuronal excitability. Furthermore, we built TBI model using *kcnt1^−/−^* mice and compared seizure susceptibility and electrophysiological properties with those of wild-type mice. We found severity-dependent seizure susceptibility in different degree of injured mice. Meanwhile, increased neuronal expression of K_Na_1.1 channel around the lesion was also observed following TBI, accompanied by a reduced firing rate of interneurons and imbalanced excitation and inhibition (E/I). Although the maximum frequency of action potential of *kcnt1^−/−^*neurons was increased, *kcnt1^−/−^* mice displayed decreased seizure susceptibility to the pentylenetetrazole (PTZ) after TBI. In *kcnt1^−/−^*mice, TBI did not significantly alter afterhyperpolarization amplitude or action-potential firing frequency in pyramidal neurons or interneurons; although the frequencies of both sEPSCs and sIPSCs increased, the E/I ratio remained unchanged. Taken together, this study suggests that enhanced K_Na_1.1 expression after TBI contributes to disputed E/I balance and seizure susceptibility, and that K_Na_1.1 might be a potential therapeutic target for reducing seizure susceptibility after TBI.

## Introduction

Traumatic brain injury (TBI) afflicts approximately ten million people throughout the world annually, contributing to increased disability and mortality *(Raymont, et al.,2010; Stein, et al.,2017)*. Post-traumatic epilepsy(PTE), a major complication, is a serious concern for neurotrauma patients and can result in the development of permanent epilepsy. The incidence of PTE varies from 4% to 53% *(Frey,2003; Zhao, et al.,2012)*, mainly depending on the injury severity *(Reid, et al.,2016)*. However, nearly one-third of patients are resistant to classical anti-seizure medications (ASMs) targeting rebalancing excitatory and inhibitory neurotransmission in the brain *(Löscher, et al.,2013)*, suggesting that TBI may induce some hitherto unknown mechanisms leading to hypersynchronous activity and seizures.

The sodium-activated potassium channel K_Na_1.1, encoded by *kcnt1 (Slo2.2)* gene, is a voltage-gated K^+^ channel that can be activated by a high cytoplasmic concentration of Na^+^. The K_Na_1.1 channel is widely expressed throughout the central nervous system (CNS) *(Rizzi, et al.,2016)*, and its functions include maintaining cellular resting membrane potential, modulating hyperpolarization following action potentials (APs), and mediating firing-rate adaptation *(Bhattacharjee and Kaczmarek,2005)*. Previous studies have reported patients carrying gain-of-function mutants of K_Na_1.1 channel are mostly with clinically ‘drug-resistant epilepsy’, such as autosomal dominant nocturnal frontal lobe epilepsy *(Heron, et al.,2012)*, malignant focal epilepsy with metastases *(Barcia, et al.,2012*; Ishii, et al.,2013), early onset epilepsy and encephalopathy *(Ohba, et al.,2015)*, and Ohtahara syndrome *(Martin, et al.,2014)*. In addition, Shore *et al*. *(Shore, et al.,2020)* found enhanced synaptic connectivity accompanied by network hyperexcitability and hyper-synchronicity after introducing a gain of function(GOF) variant of K_Na_1.1 channel in mice, which attributed primarily to reduced excitability of ‘non-fast spiking’ GABAergic neurons’ and enhanced, homotypic connectivity in both excitatory and inhibitory neurons. Although increasing studies focused on the *kcnt1*-related genetic epilepsies, there is little study relating the K_Na_1.1 channel to the acquired epilepsy. Therefore, whether K_Na_1.1 channel is involved in epileptogenesis after TBI and how K_Na_1.1 channel promotes the cortical hyperexcitability remains unknown.

In this study, we hypothesize that upregulation of K_Na_1.1 channel preferentially in inhibitory interneurons, contributes to network hyperexcitability in the perilesional cortex and enhanced seizure susceptibility after TBI. To further confirm the contributing role of the K_Na_1.1 channel upregulation to post-traumatic seizures, we induced TBI in *kcnt1* knockout *(kcnt1*^−/−^) mice and found a reduced susceptibility to seizures after TBI. The results presented here are in accord with this hypothesis.

## Results

### Behavioral Test and Tissue Loss Evaluation in Different Degrees of Injured Mice

After TBI, we assessed behavioral performance, mortality, and cortical tissue loss across different injury severities to characterize the CCI paradigms and select an injury severity for subsequent mechanistic studies (Fig. 1). Moderate and severe injury groups showed significant decreased hanging time at day 1 (Sham group: 553.2 ± 155.3 s, vs. Moderate group: 310.8 ± 241.5 s, p < 0.01; vs. Severe group: 49.9 ± 32.9 s, p < 0.001) and day 3 post-injury (Sham group: 588.3 ± 35.0 s, vs. Moderate injury group: 325.8 ± 203.3 s, p < 0.01; vs. Severe injury group: 156.2 ± 172.7 s, p < 0.001). Furthermore, severe injury group also showed significant decreased hanging time at day 7 post TBI (Sham group: 600.0 ± 0.0 s, vs. Severe injury group: 358.5 ± 239.4 s, p < 0.01). No significant difference was detected between sham and mild injury groups (Fig. 1B). Mortality within 2 weeks was also recorded to evaluate the tolerability of the different CCI paradigms. Mortality was 1/15 (6.7%) in the Sham group, 2/15 (13.3%) in the mild injury group, 3/23 (13.0%) in the moderate injury group, and 4/16 (25.0%) in the severe injury group (Fig. 1C). The single death in the Sham group was attributed to intolerance to anesthesia and was unrelated to cortical impact. In the subsequent tissue evaluation, brain slices stained by cresyl violet at 7 days after TBI were compared with the sham group, moderate and severe injury groups had significant brain tissue loss, whereas the change in the mild group was not significant (Sham group: 1.1 ± 0.81%, vs. Moderate group: 7.3 ± 2.4%, p = 0.03; vs. Severe group: 19.0 ± 2.85%, p < 0.001; Mild group: 2.9 ± 3.34%, vs. Moderate group, p = 0.04; vs. severe group, p < 0.001) (Fig. 1D, E).

**Fig. 1.**
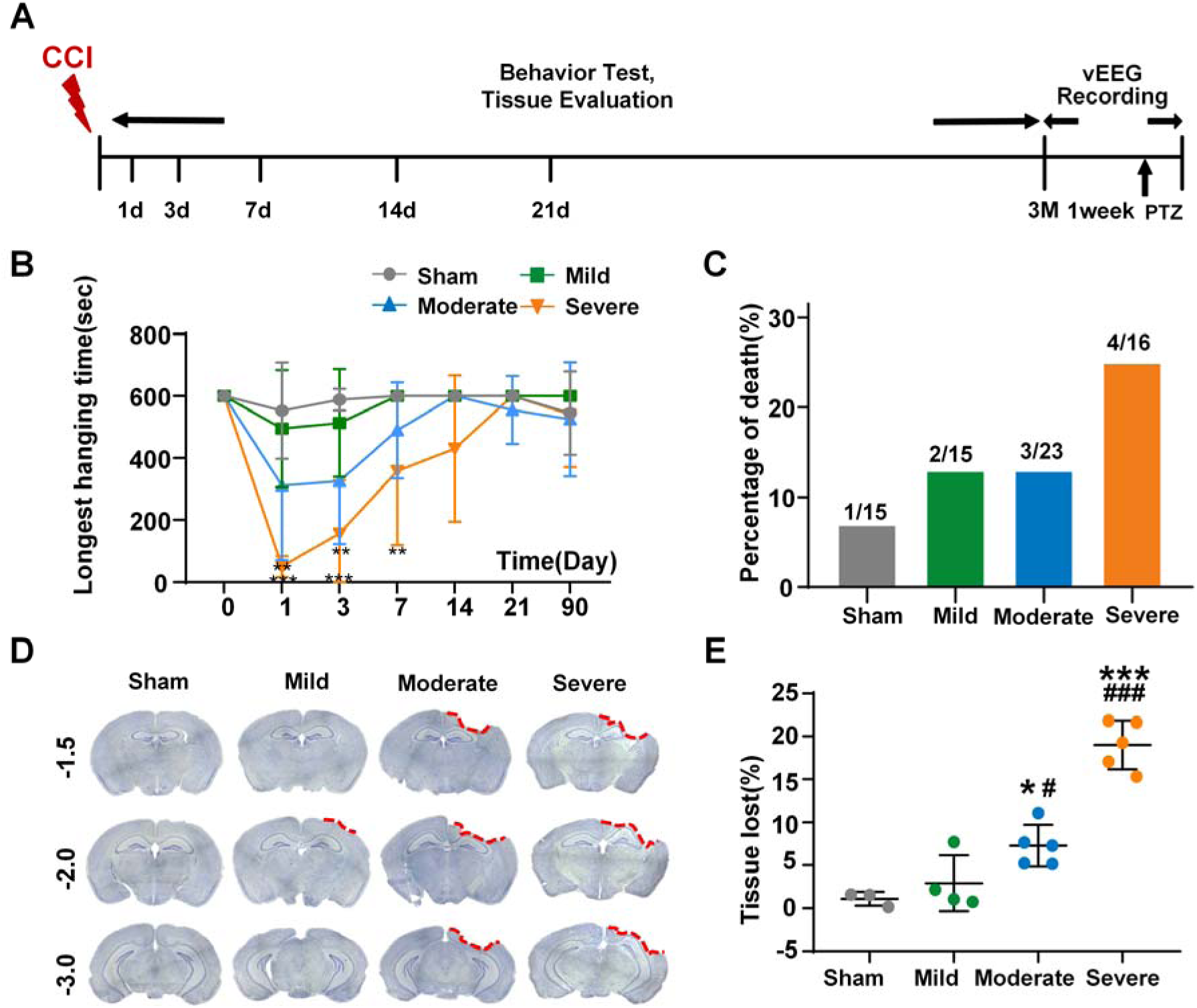
Evaluation of the degree of injury in CCI models. **A** Experimental procedure. Mice were subjected to different degrees injury with CCI models. Behavior test and tissue evaluation were performed at 1, 3, 7, 14, 21 days and 3 months post-traumatic brain injury (TBI). Skull electrodes were implanted at 11 weeks after brain damage. Long-term video-electroencephalogram (v-EEG) was monitored for one week, then followed by pentylenetetrazol (PTZ) injection. **B** Summary of behavior test in CCI models compared to sham groups at different time points. **C** Bar graph showing the percent mortality in CCI models compared to sham groups within 2 weeks. **D** Representative cresyl violet stained brain slices from three cortical regions in CCI models. **E** Summary of tissue loss in CCI models. (*p < 0.05, ***p < 0.001 vs. sham group; ^#^p < 0.05, ^###^p < 0.001 vs. mild group, Error bars indicate standard deviation).

### Severity-dependent Seizure Activity and Seizure Susceptibility after TBI in Mice

We analyzed the presence of spontaneous seizures of mice in groups with different degree of injury (Fig. 2B). While epileptiform activity was not observed in sham group, 14.3% (1/7) of mild group, 22.3% (2/9) of moderate group, and 33.3% (2/6) of severe group had at least one spontaneous seizure during consecutive 7 days of EEG recording at 3 months post-injury (Fig. 2C, left). Limited by the number of spontaneous seizures, there was no significant difference among three groups in terms of seizure frequency and duration (Fig 2C, middle and right). The spontaneous events were not systematically classified as focal versus generalized seizures; therefore, seizure type and severity were not compared across injury groups. To compare seizure susceptibility among groups, mice were challenged with a sub-convulsant dose of PTZ (35 mg/kg) at 3 months post-TBI (Fig. 2D). Compared to sham, moderate and severe groups displayed shorter latency except with the mild injury group (Sham group: 600 ± 0 s, vs. Moderate group: 141.0 ± 62.62 s, p < 0.001; vs. Severe group: 128.0 ± 40.9 s, p < 0.001; vs. Mild group: 419.5 ± 206.9 s, p = 0.61, Fig. 2E). Moreover, moderate (80%) and severe (75%) groups also exhibited generalized convulsive seizures induced by PTZ injection during the 10-min observation period, while there in no generalized convulsive seizures in sham and mild groups. The absence of generalized convulsive seizures in the mild TBI group was not interpreted as evidence of focal seizures; these mice exhibited only lower-stage seizure responses during the observation period. Specifically, the latency to generalized convulsions was significantly shorter in severe group compared to sham (Sham group: 600 ± 0 s, vs. Severe group: 322.8 ± 202.0 s, p < 0.05; Fig. 2F). In addition, the highest seizure severity scores in moderate and severe groups were significantly higher compared to sham and mild groups (Sham group: 1.0 ± 0, vs. Moderate group: 5.0 ± 1.41, p < 0.001; vs. Severe group: 5.5 ± 1.92, p < 0.001; Fig. 2G). Furthermore, more mice in severe group (2/4, 50%) died of the prolonged tonic seizure activity after PTZ injection. In sum, these results led us to utilize the moderate injury TBI model with significant motor deficit (Fig. 1B), tissue loss (Fig. 1E), lower mortality rate (Fig. 1C), and higher seizure susceptibility (Fig. 2) for the rest of our studies.

**Fig. 2.**
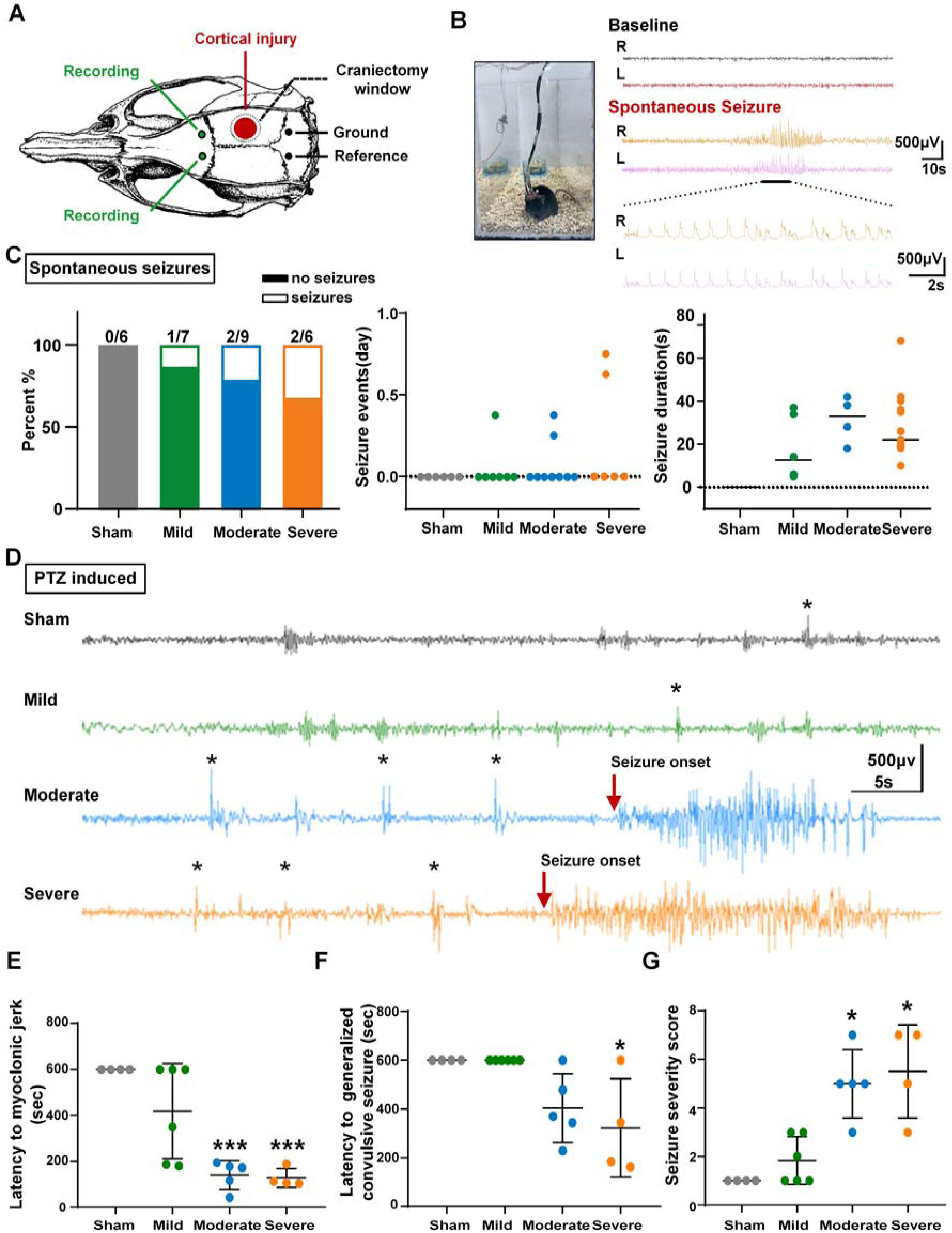
Characteristics of spontaneous seizures and pentylenetetrazol-induced seizure susceptibility in CCI models. **A** Locations of craniectomy window (black dotted circle), cortical injury (red circle), and placement of skull electrodes. **B** Long-term v-EEG monitor for 7 days on the TBI models. The epileptiform discharges of spontaneous seizure were showed in the right. **C** Evaluation of spontaneous seizures in the TBI models with different degree injury. It showed the percentage of spontaneous seizures(left), number of seizures per day (middle), and duration of seizures in TBI models. **D** Representative traces of ictal epileptiform discharges after pentylenetetrazol (PTZ) injection on the TBI models. **\***indicated interictal discharges. The red arrow indicated the onset of epileptiform discharges of generalized convulsive seizures. N = 4-6 for each group. **E** Quantification of latencies to myoclonic jerk. N = 4-6 for each group. **F** Quantification of latencies to generalized convulsive seizures. N = 4-6 for each group. **G** Quantification of seizure severity scores. N = 4-6 for each group. (*p < 0.05, ***p < 0.001. Error bars indicate standard deviation).

### Increased Expression of K_Na_1.1 Channel after TBI

K_Na_1.1 channel has been reported to be a potential therapeutic target in *KCNT1*-related epilepsies*(Milligan CJ, et al.,2014) (Burbano LE, et al.,2022)*. However, whether K_Na_1.1 channel participates in the epileptogenesis of TBI remains unknown. In this study, we collected perilesional neocortex at 1h, 6h, 12h, 1d, 3d, 7d, and 14d post-TBI and investigated spatial-temporal expression of K_Na_1.1 channel (Fig. 3A). K_Na_1.1 channel was robustly elevated in the injured ipsilateral cortex, peaking at 24 h post-TBI and maintained the high level 14 days later (Sham: 1, vs. Dpi-1h: 2.71 ± 0.70, p < 0.01; vs. Dpi-6h, 3.88 ± 1.95, p < 0.01; vs. Dpi-12h, 4.65 ± 3.07, p < 0.01; vs. Dpi-24h: 9.39 ± 5.38, p < 0.01; vs. Dpi-3d: 8.25 ± 5.53, p < 0.01; vs. Dpi-7d: 7.83 ± 3.55, p < 0.01; vs. Dpi-14d: 4.24 ± 3.33, p < 0.05, Fig. 3A). Immunofluorescence showed increased K_Na_1.1 fluorescence intensity in the perilesional cortex at dpi-14, with K_Na_1.1 signals observed in NeuN-positive cells (Fig. 3B).

**Fig. 3.**
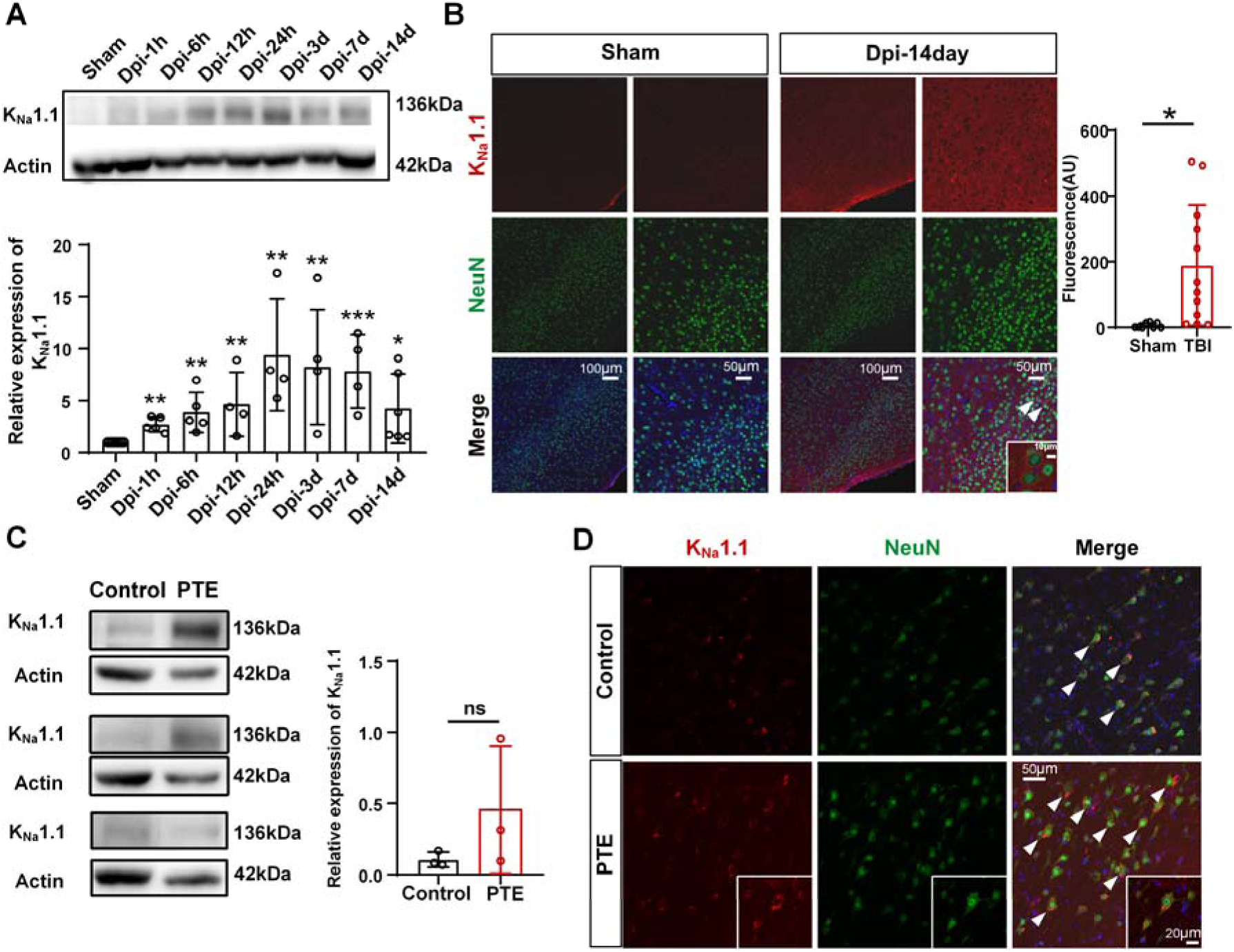
Increased K_Na_1.1 channel expression after TBI. **A** The expression of K_Na_1.1 protein at the different time points after TBI. N = 5-7 for each group. **B** At 14 days after TBI, immunofluorescence staining showed increased K_Na_1.1 fluorescence intensity in the perilesional cortex and localization of K_Na_1.1 signals in NeuN-positive cells (white arrowhead). N=3 for each group. **C** The expression of K_Na_1.1 channel on the lesion area on the neocortices from patients with PTE and patients with mesial temporal lobe epilepsy. N=3 for each group. **D** Immunofluorescence showed K_Na_1.1 signals in NeuN-positive cells, with stronger K_Na_1.1 immunoreactivity in PTE tissue than in MTLE tissue. (*p < 0.05, **p < 0.01. Error bars indicate standard deviation).

In addition to TBI mice, we also collected the lesion areas from patients with PTE and temporal neocortices from patients with mesial temporal lobe epilepsy (MTLE) as control group. The protein level of K_Na_1.1 channel was elevated in the lesion of the patient with PTE compared with that in the patient with MTLE but without no significant difference (Fig. 3C). Immunofluorescence showed K_Na_1.1 signals in NeuN-positive cells, with stronger K_Na_1.1 immunoreactivity in the PTE tissue than in the MTLE tissue (Fig. 3D).

### Decreased Expression of GAD67 and the Distribution of K_Na_1.1 in Inhibitory Interneurons in Dpi-14d Mice

Given that the protein expression of the GAD67 isoform is closely related to overall GABA levels *(Rimvall, et al.,1993)* and high expression activity of glutamate decarboxylase (GAD) is specific for mature GABAergic interneurons *(Le Magueresse and Monyer,2013)*, here we selected the GAD67 as the marker of GABAergic inhibition after TBI. Moreover, glutamatergic presynaptic structures can be labeled using antibodies against vesicular glutamate transporters (VGLUTs), especially VGLUT1 which is a subtype of VGLUTs and highly expressed in cortex *(Hussan, et al.,2022)*. Consistent with imbalance of E/I in the patch recording, western blot showed that lower GAD67 level around the lesion on the Dpi-14d mice when compared with sham group (Sham: 1 vs. Dpi-14d: 0.77 ± 0.21, p < 0.05, Fig. 4A). However, there was no significant difference in VGLUT1 level between Dpi-14d and sham groups (Fig. 4A). K_Na_1.1/VGLUT1 co-labeling showed apparent spatial overlap between the two immunofluorescence signals in sham and Dpi-14d mice (Fig. 4B). K_Na_1.1 immunoreactivity was also observed in GAD67-positive inhibitory interneurons in sham and Dpi-14d mice (Fig. 4C)

**Fig. 4.**
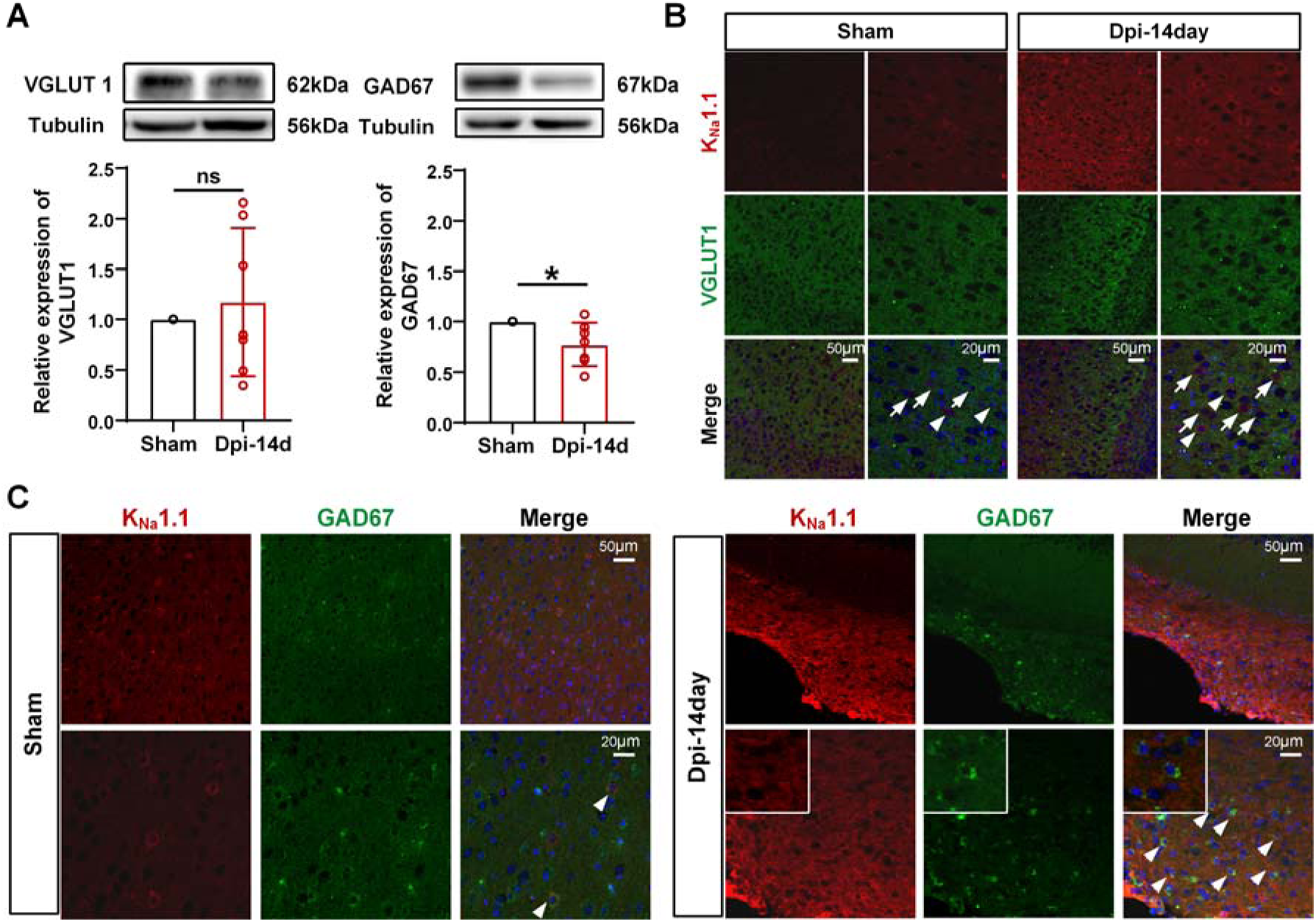
K_Na_1.1 immunoreactivity in GABAergic inhibitory interneurons in Dpi-14d mice. **A** The expression of VGLUT1 and GAD67 protein after TBI. (*p < 0.05. Error bars indicate standard deviation). **B** At Dpi-14d, immunofluorescence staining showed the distribution of K_Na_1.1 channel and VGLUT1. The white arrowheads indicated apparent spatial overlap between K_Na_1.1 and VGLUT1 signals. The white arrows indicated VGLUT1-positive profiles without apparent overlap with K_Na_1.1 signals. **C** At Dpi-14d, immunofluorescence staining showed the distribution of K_Na_1.1 channel and GAD67 on neurons. The white arrowheads indicated the neurons with co-localized K_Na_1.1 channel and GAD67. Panels B and C were assessed qualitatively; no quantitative co-localization analysis was performed.

### Imbalance of Glutamatergic and GABAergic Neurons in Perilesional Cortex in Moderate TBI Mice

Previous studies have demonstrated the development of cortical hyperexcitability during the subacute period after TBI. Nichols et al. reported epileptiform activity within 14 days after CCI in juvenile rats *(Nichols, et al.,2015)*, while increased perilesional cortical neuronal firing at 14 days post-injury has also been observed in a mouse CCI model *(Ping X, et al.,2016)*. Therefore, we selected 14 days post-TBI as a subacute time point to assess neuronal intrinsic excitability and synaptic activity. For the Dpi-14d mice, we assessed neuronal subtype-specific AP characteristics and synaptic activity by performing whole-cell patch-clamp recordings of neurons in the perilesional neocortex of post-injury moderate TBI mice (Fig. 5A). Compared with the sham group, maximum AP frequency (Sham: 45.3 ± 9.41 Hz vs. TBI: 32.2 ± 8.34 Hz, p < 0.05, Fig. 5C, 5D) and afterhyperpolarization (Sham: 6.4 ± 2.90 mV vs. TBI: 10.8 ± 3.31 mV, p < 0.05, Fig. 5C, 5D) of interneurons (INs) showed significant differences. However, there was no difference on these electrophysiological parameters of pyramidal neurons (PNs) (Fig. 5B, 5D). We calculated the resting membrane potential between Sham and TBI groups and found there was no significant difference in both PNs and INs (data not shown). Next, we evaluated spontaneous excitatory and inhibitory post-synaptic currents (sEPSC and sIPSC) in TBI and sham groups. The sEPSC frequency of PNs in the TBI group was significantly higher than that of the sham group (Sham: 2.14 ± 1.33 Hz vs. TBI: 4.57 ± 3.94 Hz, p < 0.05, Fig. 5E, 5F, left), while the sIPSC frequency was markedly reduced (Sham: 1.74 ± 1.18 Hz vs TBI: 0.54 ± 0.54 Hz, p < 0.001, Fig. 5E, 5F, middle), and excitation and inhibition (E/I) ratio was significantly increased (Sham: 0.38 ± 0.30, vs TBI: 0.74 ± 0.29, p < 0.001, Fig. 5F, right). This data indicates that the excitatory driving force of PNs in the cortex increased, while the inhibitory driving force decreased, hence the overall excitability was enhanced in the perilesional neocortex after TBI. The synaptic activity of INs showed no difference between sham and TBI groups (data not shown).

**Fig. 5.**
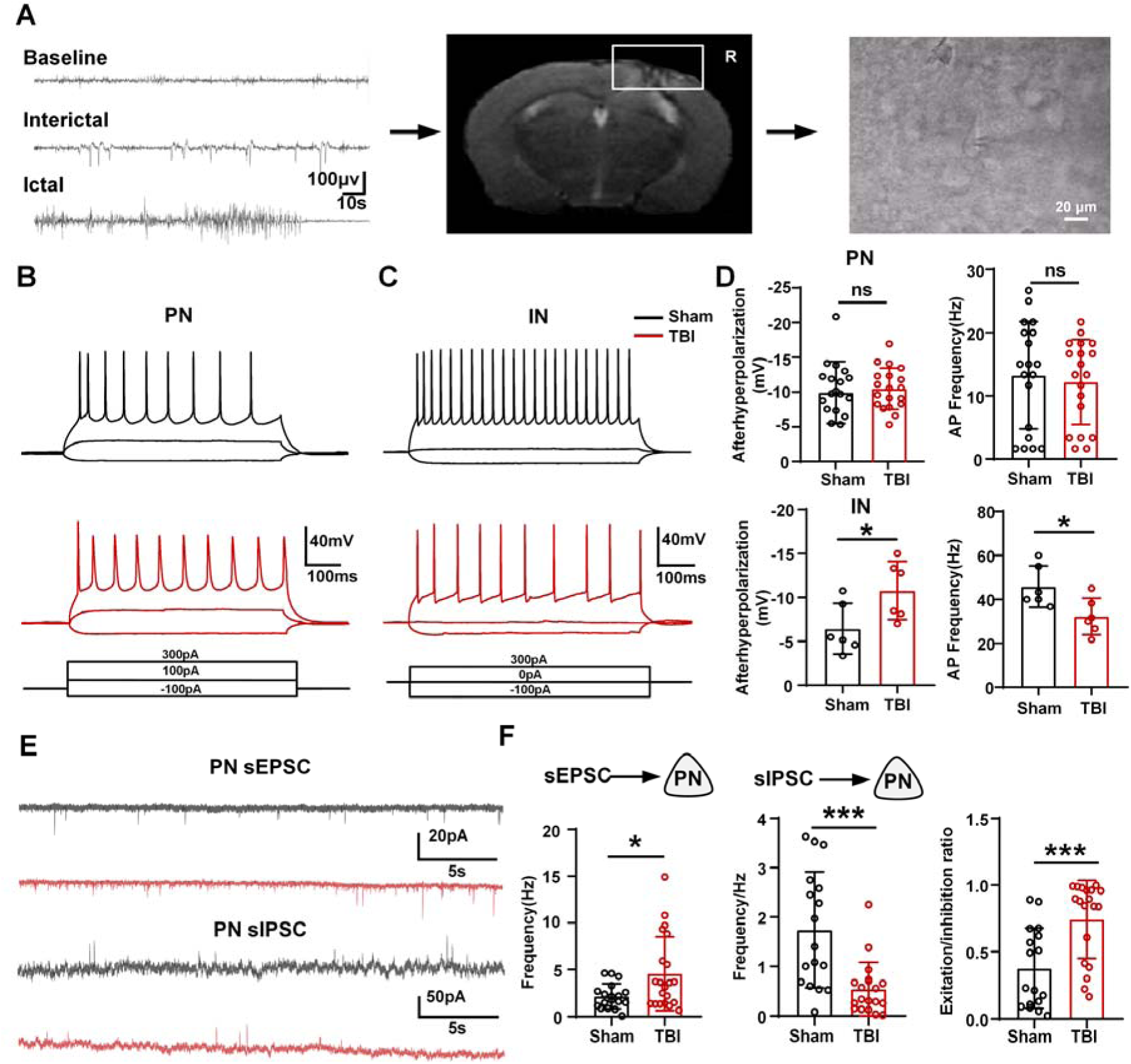
Altered electrophysiological characteristics of perilesional TBI neurons. **A** In vivo video-EEG recordings showed interictal and ictal discharges in TBI mice (left). Neurons in layer II/III of perilesional cortex were selected for ex vivo whole-cell patch-clamp recordings in acute cortical slices (middle and right). **B** Representative traces and action potential (AP) firing of PNs from sham and TBI groups with – 100, 100, +300 pA current injection. **C** Representative traces and AP firing of INs from sham and TBI groups with –100, 0, +300 pA current injection. **D** Summary of afterhyperpolarization and AP frequency in PNs and INs. PNs: N = 18, 21 for sham and TBI groups. INs: N = 6 for sham and TBI groups. **E** Representative traces of spontaneous excitatory postsynaptic currents (sEPSC) and spontaneous inhibitory postsynaptic currents (sIPSC) of PNs (Black, sham group; red, TBI group). N = 18, 21 for sham and TBI groups. **F** Quantitative summary of sEPSC and sIPSC frequency and excitation/inhibition ratio of PNs. N = 18, 21 for sham and TBI groups. (*p < 0.05, ***p < 0.001. Error bars indicate standard deviation).

### Improved Behavior and Decreased Seizure Susceptibility on *kcnt1^−/−^* Mice after TBI

To figure out the functional role of K_Na_1.1 channel in TBI, we created a loss of *kcnt1* mouse line whose mutation site lies adjacent to the pore-formation domain of K_Na_1.1 channel (Fig. 6A). We confirmed cohorts of wild-type and homozygous *(kcnt1^−/−^)* littermates using tail PCR (Fig. 6B). Genetic transmission of both alleles was normal, and no selective lethality was seen in either mutant genotype at any age. Western blot analysis of protein fractions isolated from the cortices of WT and *kcnt1^−/−^* mice showed that K_Na_1.1 channel was detected only in the WT mice, while K_Na_1.1 expression was significantly higher than WT group at 7 days after TBI (Fig. 6C). Extracted brains from 10 to 12-week-old male *kcnt1^−/−^*mice appeared similar in morphology and weight to their WT littermates (Fig. 6D), suggesting that loss of the *kcnt1* does not grossly alter brain development.

**Fig. 6.**
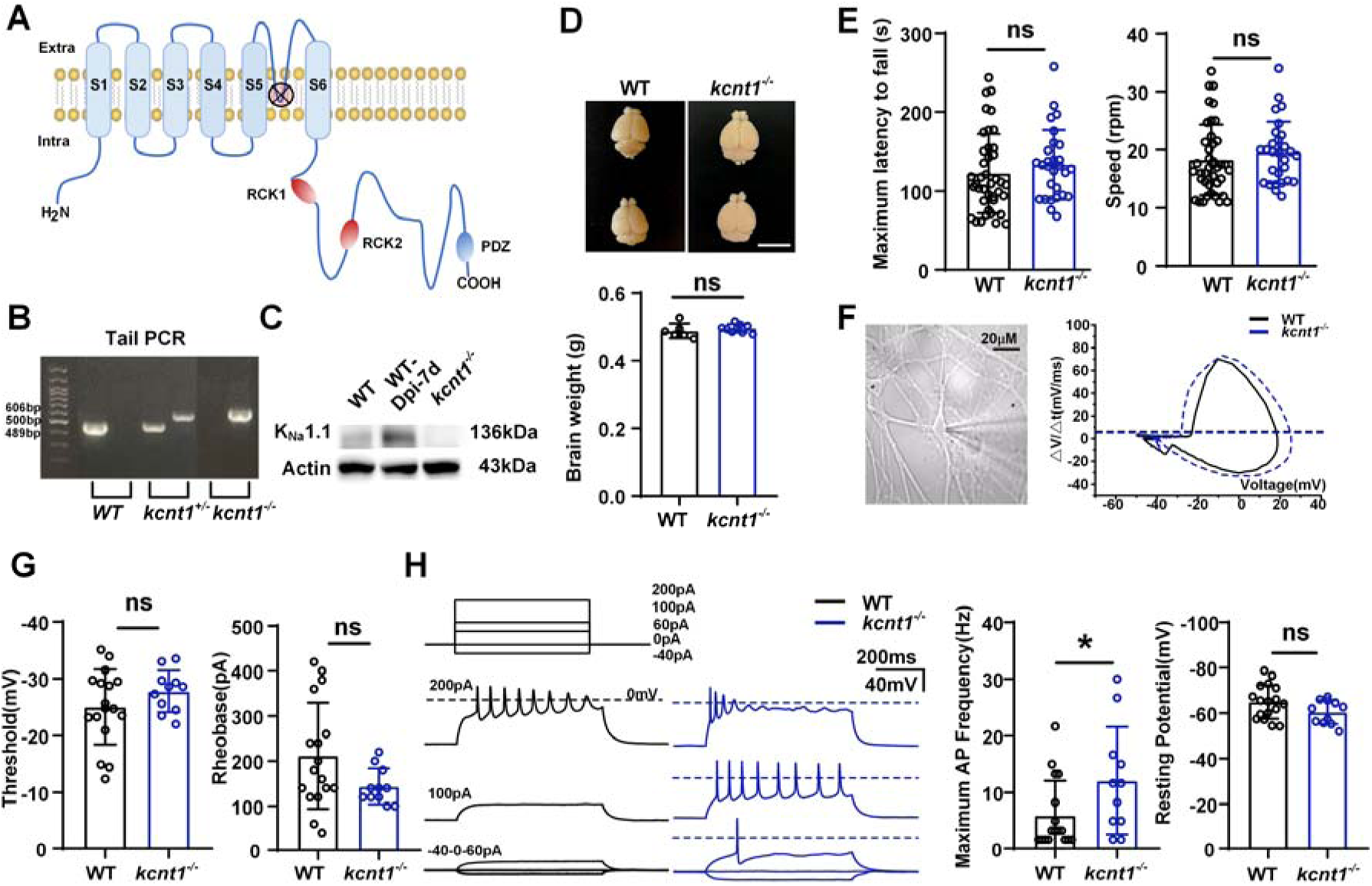
Evaluation of homozygous *kcnt1*^-/-^ mice and age-matched wild type controls. **A** Diagram of K_Na_1.1 protein in the *kcnt1*^−/−^ mice showed corresponding mistranslated site in its pore-formation domain. **B** PCR verification of *kcnt1* exon 3-11 deletion. The predicted amplicons for WT and the exon 3-11 deleted mutant were 489 bp and 606 bp, respectively. The corresponding sample from the WT, *kcnt1*^+/-^, and *kcnt1^−/−^*. **C** Western blots showed K_Na_1.1 expression from the WT, WT-TBI (for a positive control), and *kcnt1^−/−^* mice. **D** Representative image showed similar gross morphology of brains dissected from 10 to 12-week-old male WT, *kcnt1^−/−^* mice. N = 7 in WT and *kcnt1*^−/−^ groups, respectively. Scar bar represented the length of 1cm. **E** Evaluation of motor task in *kcnt1*^−/−^ and WT mice (WT, black; *kcnt1^−/−^*, blue). N = 40, 28 in WT and *kcnt1*^−/−^ groups, respectively. **F** Representative diagram showed cultured primary cortical neuron from *kcnt1*^−/−^ mice. dV/dt is plotted as function of membrane voltage for the action potentials (APs) (WT, black; *kcnt1^−/−^*, blue). Horizontal dotted lines correspond to the dV/dt value that is 10% of peak dV/dt for a given cell. **G** Comparisons of AP threshold and rheobase of cultured primary cortical neuron from WT and *kcnt1*^−/−^ mice. N= 17, 11 in WT and *kcnt1*^−/−^ groups, respectively. **H** –40, 0, 60, 100, and 200 pA current injections (0.6 s) were used to elicit firing in WT and *kcnt1^−/−^* (blue) neurons from a holding potential of −70 mV. AP frequency at each current step was calculated as the total number of action potentials during the entire 600-ms pulse divided by 0.6 s. For each neuron, the maximum frequency across the tested depolarizing current steps was used for the genotype comparison. Comparisons of maximum AP frequency and resting membrane potential of cultured primary cortical neuron from *kcnt1*^−/−^ and WT mice. N= 17, 11 in WT and *kcnt1*^−/−^ groups, respectively. (*p < 0.05. Error bars indicate standard deviation).

In addition, motor skill was tested by measuring the falling time and speed which animals were able to maintain balance on a rotarod with repeated trials. There was no significant difference in the running time or speed between WT and *kcnt1^−/−^* mice (Fig. 6E). Furthermore, to characterize the baseline electrophysiological phenotype associated with K_Na_1.1 deletion, we compared the intrinsic membrane properties of cultured primary cortical neurons between *kcnt1^−/−^* and WT mice (Fig. 6F-H). Although there was no significant difference in terms of AP threshold, rheobase, and resting membrane potential between groups, the maximum frequency of AP on *kcnt1^−/−^*neurons was increased when compared with WT neurons (WT: 6.0 ± 6.13, vs. *kcnt1^−/−^*: 12.1 ± 9.46, p < 0.05).

### Decreased seizure susceptibility in *kcnt1^−/−^* mice after TBI

We named these groups as Sham (WT mice), TBI (WT mice undergoing TBI), *kcnt1*^−/−^-Sham*(kcnt1*^−/−^ mice without TBI), and *kcnt1*^−/−^-TBI *(kcnt1*^−/−^ mice undergoing TBI). We evaluated motor performance among the four groups using the rotarod test. Compared with the Sham group, the TBI group showed significantly lower maximum latency to fall and speed at dpi-1. The *kcnt1*^−/−^-Sham group showed significantly higher maximum latency to fall and speed than the TBI group at dpi-3 and dpi-7. At dpi-14, the *kcnt1*^−/−^-TBI group showed significantly higher maximum latency to fall and speed than the Sham group, and its speed was also significantly higher than that of the TBI group (Fig. 7B).

**Fig. 7.**
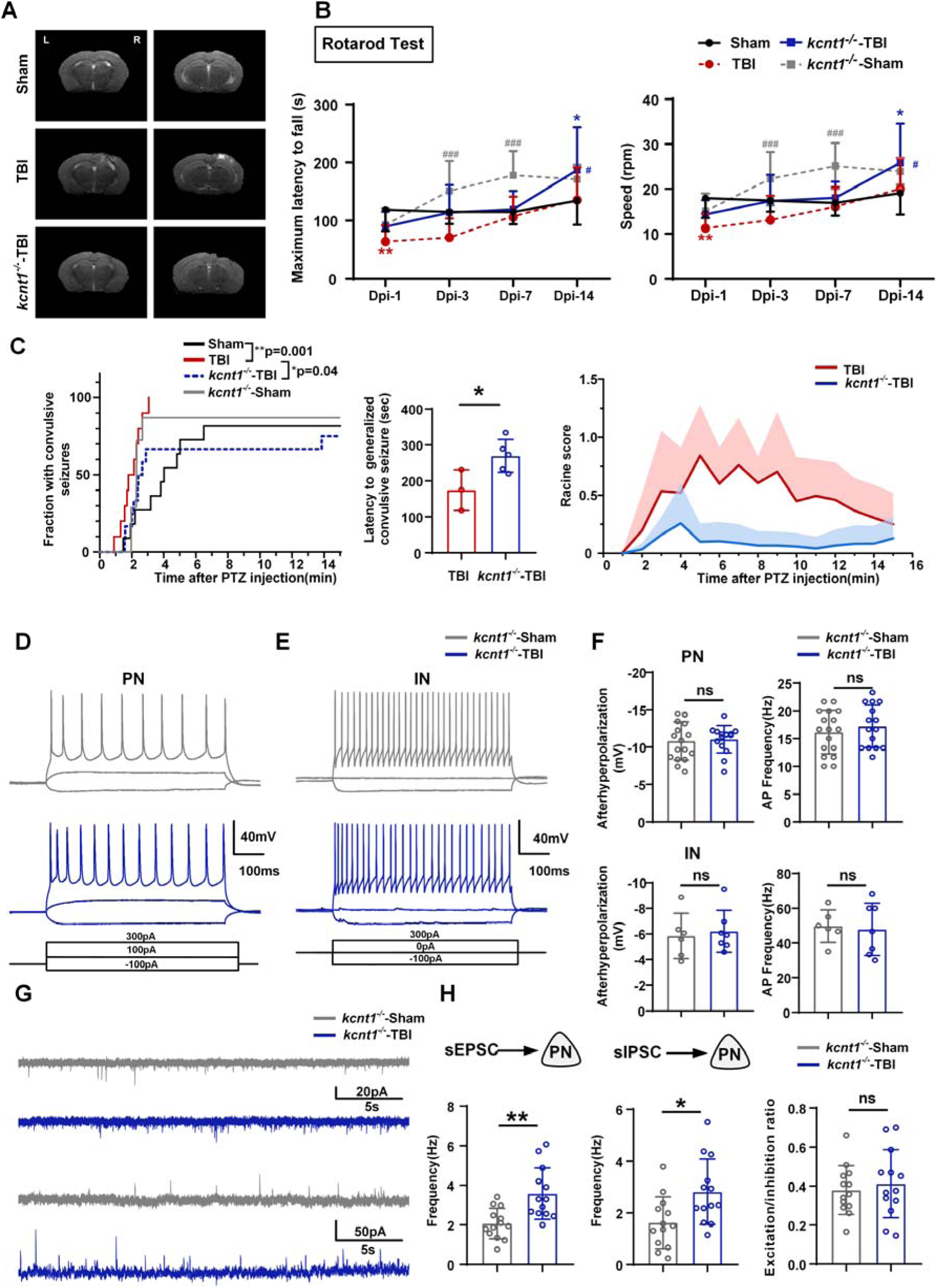
Decreased seizure susceptibility in homozygous *kcnt1*^-/-^ mice after TBI. **A** Representative MRI image showed both TBI and *kcnt1^−/−^* –TBI mice appeared lesions in the cortex after TBI. **B** The motor performance of WT and *kcnt1^−/−^* mice with or without TBI was evaluated using the accelerating rotarod test over successive trials. *p < 0.05, **p < 0.01, ***p < 0.001 vs. Sham group; #p < 0.05, ##p < 0.01 vs. TBI group. Initial N = 11, 8, 17, and 16 for Sham, *kcnt1^−/−^*-Sham, TBI, and *kcnt1^−/−^*-TBI groups, respectively. **C** The latency to seizure, including myoclonic jerk and generalized clonus, and distribution of Racine scores after injection of PTZ. N = 11,9,7,12 for Sham, TBI, *kcnt1^−/−^*-sham, and *kcnt1^−/−^*-TBI groups, respectively. **D** Representative traces and AP firing of PNs from *kcnt1^−/−^*-Sham and *kcnt1^−/−^*-TBI group with –100, 100, +300 pA current injection. **E** Representative traces and AP firing of INs from *kcnt1^−/−^*-Sham and *kcnt1^−/−^*-TBI group with –100, 0, +300 pA current injection. **F** Summary of afterhyperpolarization and AP frequency in PNs and INs. PNs: N = 17, 15 for *kcnt1^−/−^*-Sham and *kcnt1^−/−^*-TBI group. INs: N = 6, 7 for *kcnt1^−/−^*-Sham and *kcnt1^−/−^*-TBI group. **G** Representative traces of sEPSC and sIPSC of PNs (Grey, *kcnt1^−/−^*-Sham group; blue, *kcnt1^−/−^*-TBI group). N =13 for *kcnt1^−/−^*-Sham and *kcnt1^−/−^*-TBI group. **H** Quantitative summary of sEPSC and sIPSC frequency and excitation/inhibition ratio of PNs. N =13 for *kcnt1^−/−^*-Sham and *kcnt1^−/−^*-TBI group. (*p < 0.05,**p < 0.01, ***p < 0.001. Error bars indicate standard deviation).

We next tested the effects of deletion of the rodent *kcnt1* gene on seizure susceptibility by administration of 35 mg/kg PTZ at 14 days after TBI. In all animals, seizures were monitored within 15 minutes following PTZ administration. There was no significant difference in seizure susceptibility between *kcnt1*^−/−^-sham and WT-sham groups. However, latency to convulsive seizures (including myoclonic jerk and generalized clonus) following PTZ showed longer latencies in *kcnt1*^−/−^-TBI group compared with TBI group (p = 0.04, log rank test, Fig. 7C, left). Moreover, there was a significant difference in latency to generalized convulsive seizure, which were 174.7 ± 56.5 s and 269.8 ± 45.65 s for TBI and *kcnt1*^−/−^-TBI group respectively (Fig. 7C, middle). However, compared with *kcnt1*^−/−^-TBI group, we observed longer seizure duration and more severe seizures in TBI group as evidenced by significant differences in the distribution of Racine scores *(kcnt1*^−/−^-TBI vs TBI, Kruskal-Wallis test, p < 0.001, Fig. 7C, right).

We further examined the electrophysiological properties of perilesional cortical neurons in *kcnt1^−/−^*-Sham and *kcnt1^−/−^*-TBI mice at 14 days after TBI. No significant differences in afterhyperpolarization amplitude or action-potential firing frequency were detected in either PNs or INs between the two groups (Fig. 7D-F). The frequencies of both sEPSCs and sIPSCs were significantly increased in PNs from *kcnt1^−/−^*-TBI mice compared with *kcnt1^−/−^*-Sham mice, whereas the E/I ratio did not differ significantly between the groups (Fig. 7G, H).

## Discussion

In this study, we firstly evaluated the properties of TBI mice with different degrees of injury and found its severity-dependent behavior deficit, tissue damage and seizure activities. Next, we selected the moderate TBI mice as study subjects and found enhanced seizure susceptibility, increased neuronal excitability including reduced firing rate of interneurons and imbalanced excitation and inhibition (E/I) in the moderate TBI model combining EEG recordings with patch clamp recordings. Increased expression of K_Na_1.1 channel in TBI mice, preferentially in inhibitory interneurons, may underly in the disputed E/I and increased seizure susceptibility. Meanwhile, a rise in K_Na_1.1 channel was also found in patients with PTE compared with controls from patients with MTLE. To determine the functional role of K_Na_1.1 channel after TBI, *kcnt1^−/−^*mice undergoing TBI displayed improved motor learning skill and decreased seizure susceptibility to the PTZ. Although there has been a long-held view that GOF of K_Na_1.1 channel in the brain plays a pivotal role in the *kcnt1*-related epilepsy, we demonstrate for the first time that enhanced K_Na_1.1 channel after TBI with changed distribution on different subtypes of neurons, which contributes to impaired cortical inhibition, E/I imbalance, and enhanced seizure susceptibility during the subacute post-traumatic period. In doing so, this research was able to reveal a critical target for prevention and treatment during the critical subacute post-traumatic period.

### Severity-dependent deficits prompt moderate CCI model a suitable study subject

Well-designed preclinical PTE models can help to evaluate mechanisms involved in PTE which cannot be determined in human patients. The animal model of TBI mentioned here mimics focal brain trauma, which is prone to the development of PTE compared with other models *(Hunt, et al.,2009)*. Similar with human studies, one critical risk factor of PTE on the CCI model is the injury severity *(Curia, et al.,2011)*. In this study, we utilized sham and three severity levels of CCI mice and found that neurological function deficits, cortical tissue loss and increased susceptibility to PTZ were severity-dependent after TBI. Indeed, the more severe the injury, the more pronounced features observed. Considering the combination of measurable neurological deficits, cortical tissue loss, increased seizure susceptibility, and lower mortality than the severe injury group, we selected the moderate CCI model for subsequent mechanistic experiments.

One study found that the incidence of spontaneous seizures was only 9% in mice followed up for 9 months after TBI, with fairly infrequent seizures (0.2-0.3 seizures/day) *(Bolkvadze and Pitkänen,2012)*. Inconsistent with the previous study, spontaneous seizures in our study are only about 0.3 seizures/day. Capturing these sporadic seizures is challenging in terms of animal numbers and long-term recording, which hinders high-throughput studies. Although the clinical relevance of enhanced susceptibility to PTZ remains controversial, evidence suggests that PTZ-evoked hyperexcitability is associated with the potential to develop to spontaneous seizures *(Blanco, et al.,2009; Rattka, et al.,2011)*. Here, we employed PTZ as a measure to evaluate the seizure susceptibility by injecting a sub-convulsive dose into mice after injury. Subsequent EEG monitoring revealed that the response of mice with moderate or severe injuries to proconvulsant PTZ was significantly faster and stronger than that observed in sham and mild injury groups, indicating that the increase in seizure susceptibility was proportional to the degree of damage, consistent with clinical patients *(Annegers, et al.,1998)*.

### Imbalance of E/I in perilesional neocortex might contribute to epileptogenesis

Previous studies indicate that post-traumatic cortical hyperexcitability can emerge during the early to subacute period after injury *(Nichols, et al.,2015) (Ping X, et al.,2016)*. However, its temporal progression may vary depending on species, developmental stage, and injury model. Accordingly, the 14-day time point in the present study was used to characterize subacute neuronal and network alterations rather than to define the onset of PTE. Although partial seizure is the dominant seizure type after brain trauma, few studies have definitively determined which cortical regions are selectively predisposed to epileptogenic vulnerability. Human postsurgical pathology revealed iron deposition, nonspecific gliosis and fibrotic meningo-cortical scarring occurred adjacent to the injured cortex, which has been identified as a potential epileptic zone *(Clifton, et al.,1981; Marks, et al.,1995)*. However, there are few corresponding functional studies that evaluated the consistency between epileptogenic area and perilesional neocortex. In fact, the excitability of neural networks and hyperactivity have changed significantly before post-traumatic epileptogenic foci can be identified.

However, most studies mainly focus on the macroscopic epilepsy expression with neglect on microscopic detection in the epileptogenic mechanisms. Thus, Farrell et al. *(Farrell, et al.,2019)* highlighted that micro-macro disconnection should be noticed in uncovering the detailed epileptogenic mechanisms. Here, inspired by enhanced neuronal activities after TBI using EEG recording, we performed patch clamp recording on the perilesional neocortex as well to resolve this disconnection. Our results indicated that the intrinsic membrane properties of PNs remain unchanged after CCI. Contrary to PNs, AP characteristics were altered in inhibitory INs, possibly reflecting unique TBI-induced changes to inhibitory INs. At the synaptic level, the frequency of sEPSC and sIPSC in PNs were markedly altered following CCI, and E/I ratio was significantly increased, indicating that the overall excitability was enhanced in the perilesional neocortex after TBI. Overall, our results suggested that specific cortical neuronal microcircuits, such as increased glutamatergic signaling and reduced GABAergic inhibition, may initiate and facilitate the spread of epileptiform activity in the perilesional neocortex after TBI. Consisted with Cantu et al. *(Cantu, et al.,2015)*, we speculate that hyperexcited network at macroscale is due to imbalance between E/I and the loss of GABAergic control at microscale.

### The role of enhanced K_Na_1.1 channel in post-traumatic cortical hyperexcitability and seizure susceptibility

Since *kcnt1* was confirmed as an epilepsy-associated gene in 2012, GOF mutations have been reported to be closely related to “refractory epilepsy” in infants. Increasing studies expand current understanding of the epileptogenesis mechanisms of *kcnt1*-associated epilepsy from remodeling heterologous channel with pathogenic variants to initial characterization of mouse models. However, our study firstly reported that enhanced K_Na_1.1 channel contributes to post-traumatic cortical hyperexcitability and increased seizure susceptibility in an acquired brain injury model. Herein, we provided the evidence that expression of K_Na_1.1 channels was elevated in neurons from the perilesional cortex. Qualitative immunofluorescence images showed K_Na_1.1 immunoreactivity in GAD67-positive inhibitory interneurons in the perilesional cortex. However, the role of enhanced K_Na_1.1 channel in epileptogenesis of TBI still remains unclear. Consistent with previous studies about the detailed mechanisms of *kcnt1*-related epilepsy, one plausible explanation of enhanced K_Na_1.1 channels after TBI is that the variants in *kcnt1* may result in an imbalance of its expression in excitatory and inhibitory neurons *(Tang, et al.,2016)*. In other words, when K_Na_1.1 channel is over-expressed in inhibitory neurons, it will increase overall neuronal excitability by reducing the excitability of inhibitory neurons *(Bhattacharjee, et al.,2002; Joshi, et al.,2012)*. In addition, changing synaptic connectivity caused by GOF in K_Na_1.1 channels possibly participates in the epileptogenic process. Shore and colleagues introduced a GOF mutation of K_Na_ channel (KCNT1-Y777H) into mice and, interestingly, they observed a cell-type-specific difference on K_Na_ currents. They further provided evidence of synaptic rewiring, including homotypic synaptic connectivity *(Shore, et al.,2020)*. Another reasonable explanation for these conflicting findings is that there is an abnormal distribution of K_Na_1.1 channel at the subcellular level. Our previous study found that activation of K_Na_1.1 channel has a braking effect on the initiation of Aps *(Martinez-Espinosa, et al.,2015)*. When K_Na_1.1 channel was overexpressed in non-initiating regions of actions potentials, such as the peripheral axon, then the braking effect was reduced and neuronal excitability was increased. However, these hypotheses all need further investigation.

To confirm the role of K_Na_1.1 channels in TBI, we produced *kcnt1^−/−^* mice by deleting the exon 3–11 region of the *kcnt1* gene and compared behavior tests and neuronal properties between WT and *kcnt1^−/−^* mice. Although WT and *kcnt1^−/−^* mice displayed similar motor activity in rotarod test, neurons from *kcnt1^−/−^*mice were found with increased AP firing rate. As for seizure susceptibility by administration of PTZ, there was no significant difference in latency to seizures between sham and *kcnt1^−/−^*-sham mice. However, *kcnt1*^−/−^-TBI group was improved in procedural learning and reduced in the seizure susceptibility than TBI group at dpi-14. Furthermore, we performed whole-cell patch clamp recordings on the perilesional neocortex of *kcnt1^−/−^*-Sham and *kcnt1^−/−^*-TBI mice at dpi-14. There was no significant difference in afterhyperpolarization or AP firing frequency of either PNs or INs between the two groups. Although the frequencies of both sEPSC and sIPSC in PNs were significantly increased after TBI in *kcnt1^−/−^* mice, the E/I ratio showed no significant difference. In contrast to the reduced firing rate of INs and imbalanced E/I observed in WT-TBI mice, these alterations were not observed in *kcnt1^−/−^*-TBI mice. These results suggested that K_Na_1.1 channel might contribute to impaired inhibitory neuronal excitability and imbalanced E/I after TBI. In our study, consistent with previous studies, we found that the maximum frequency of AP of *kcnt1^−/−^* neurons was increased when compared with WT neurons, indicating that K_Na_1.1 might be a protective modulator in a physiological manner. However, pathological redistribution of K_Na_1.1 channels may contribute to impaired inhibitory neuronal excitability, E/I imbalance, and increased seizure susceptibility after TBI. In contrast to genetic KCNT1-related epilepsy, our findings suggest that altered K_Na_1.1 distribution following TBI contributes to post-traumatic E/I imbalance, network hyperexcitability, and increased seizure susceptibility.

### Outlooks and limitations

This study had several inevitable limitations. Firstly, in terms of the exact role of upregulation of K_Na_1.1 channel after TBI, we admitted that we could not solve it undisputedly in this study. When seizure occurs, hypersynchronous electrical activity calls for a sudden influx in Na^+^ and activates K_Na_ 1.1 channel which is sensitive to Na^+^. Byers et al. reported that K_Na_ 1.1 channel, activated by neuronal overexcitation, contribute to a protective threshold to suppress the induction of seizure-like activity not only in established genetic but also pharmacologically primed seizure models *(Byers, et al.,2021)*. Although we found the distribution difference in K_Na_1.1 channel between PNs and INs after TBI, we could not exclude other factors which might regulate the function of K_Na_1.1 channel, such as factors linked with K_Na_ channels by modulate intracellular Na^+^ or interact with K_Na_ channels via effecting its Na^+^ sensitivity *(Aoki, et al.,2018; Tamsett, et al.,2009)*. Secondly, there is the absence of reversing verification with established *kcnt1* over-expression animals undergoing TBI. Given that neurons are very sensitive to expressed levels of K_Na_ 1.1 channel, it is hard to titrate the right amount of K_Na_ 1.1 channel expression. In addition, post-TBI tissue loss was not quantitatively compared between WT and *kcnt1^−/−^* mice. Therefore, we cannot determine whether the reduced PTZ-induced seizure susceptibility in *kcnt1^−/−^* mice reflects reduced tissue injury, altered neuronal intrinsic properties, or both. Thirdly, molecular and electrophysiological changes were examined within 14 days after TBI, whereas spontaneous seizures were monitored for only 7 days at 3 months post-injury. This limited v-EEG window may have missed infrequent events, and PTZ challenge reflects induced seizure susceptibility rather than spontaneous PTE. Therefore, the early alterations observed here cannot establish a causal relationship with chronic PTE, which requires long-term or repeated v-EEG monitoring in WT and *kcnt1^−/−^*mice. Fourthly, NeuN immunoreactivity may be altered after TBI. Therefore, NeuN staining was used only to determine the neuronal localization of K_Na_1.1 rather than to quantify neuronal survival. Future studies should combine Nissl staining or neuronal degeneration markers to assess neuronal loss. The specificity of K_Na_1.1 immunofluorescence staining was not directly validated using *kcnt1^−/−^*brain sections. Similarly, reduced GAD67 protein levels alone do not establish a reduction in inhibitory neuron number. Furthermore, miniature inhibitory postsynaptic currents (mIPSCs) were not recorded, limiting our ability to distinguish changes in action-potential-dependent network activity from alterations in quantal inhibitory synaptic transmission. The present findings support impaired inhibitory signaling after TBI but do not establish a specific presynaptic mechanism or inhibitory neuron loss. Fifthly, the size of our patient cohort was relatively small. In the future, a larger cohort of patients should be recruited to confirm the change and role of K_Na_1.1 in patients with PTE, which might provide a novel direction for treatment.

## Materials and Methods

### Animals

A null mutant-*kcnt1^−/−^* mice were obtained that were initially generated on a C57BL/6J background by deleting the exon 3-11 of *kcnt1* gene sequence (NCBI: 227632) and homologous recombing in murine embryonic stem cells. *kcnt1^−/−^*and wild-type C57BL/6 background strain (Vital River Laboratory Animal Technology Co., Ltd, Beijing, China) were subsequently backcrossed at least 2 generations prior to testing. Adult male WT and *kcnt1^−/−^* male mice (8-10 weeks old) were housed in an animal facility with suitable temperature and humidity, a 12h light-dark cycle, and free access to food and water. All procedures were approved by the Animal Care and Use Committee of Capital Medical University in accordance with the Guide for the Care and Use of Laboratory Animals.

### Human Tissue

The studies involving human participants were reviewed and approved by the Medical Ethics Committee of Tiantan Hospital, Capital Medical University. Written informed consent to participate in this study was provided by the participants’ legal guardian/next of kin.

### TBI Models

For TBI models, mice were anesthetized with 3% isoflurane in 67% N_2_/30% O_2_ mixture, placed in a stereotaxic frame, and maintained with 1.5% isoflurane. Brain injury was delivered using controlled cortical impact (CCI) equipment (Leica 9969S; Wetzlar, Germany). We performed a craniectomy (4 mm diameter) from the bregma over the right cortex region (center: AP –2.00 mm, RL +2.00 mm). CCI equipment was mounted on the right stereotaxic arm at a 15° angle from vertical. The diameter of the impactor was 3 mm (flat tip). Impact depths were 0.5 mm (mild), 1.0 mm (moderate), or 1.5 mm (severe), delivered with a velocity of 3.0 m/s and 100 ms duration. Sham-injured mice received only craniectomy and were not subjected to CCI impact. The timeline diagram depicts timing of the experiments through the TBI protocol and corresponding experiments (Fig. 1A).

### Primary Cortical Neuron Culture

For primary neuron cultures, cortices from WT and *kcnt1^−/−^*mice aged postnatal day 0-1 and of either sex were dissected in cold phosphate-buffered saline (PBS). The tissue was then cut into pieces and digested with 0.05% trypsin (T1300; Solarbio, Beijing, China) and DNAase (01086251; Thermo Fisher Scientific, Waltham, MA, USA) for 10-20 min. Digestion was terminated by adding Dulbecco’s Modified Eagle Medium (DMEM; 11995-065; Gibco, Amarillo, TX, USA) supplemented with 10% fetal bovine serum (10099-141C, Gibco), 10% horse serum (04-004-1A; Biological Industries, Beit Haemek, Israel), and 50× penicillin/streptomycin (15140-122, Gibco). After filtering the cell suspension with 70 μm mesh, it was centrifuged at 1000 rpm, dispersed, and resuspended with DMEM. Next, cells were mechanically counted and added at densities of 1 × 10^5^ and 1 × 10^6^ cells/well to poly-D-lysine pre-coated 24-well or 6-well plates, respectively. After 4h incubation to allow for cell attachment, DMEM was replaced with NBA Plus [Neurobasal-A medium (10888-022, Gibco) supplemented with 1× GlutaMAX (35050-061, Gibco), 1× B27 supplement (17504-044, Gibco), and 100× penicillin/streptomycin]. Cells were treated with cytarabine (5 μM) and cultured for 3-5 days to inhibit the proliferation of gliocytes. Every 2 days, 50% of the media was replaced with fresh NBA Plus.

### Behavioral Tests

In the behavioral 75% alcohol was prepared to eliminate substances and smell that left in machine between two mice to be tested. For the rotarod test, mice were pre-trained on 3 consecutive days on the rotarod rotating for 3 times with the interval of 5 min. Mice were then tested at the initial speed of 4 rpm, accelerating speed ranging from 4 to 30 rpm within 5 min. Time and speed spent on the rotating rod until falling off were recorded for each performance. Each test including three repetitions with an inter-trial interval of 5 min and the means from these three runs were analyzed for each mouse.

For the hanging test to evaluate the movement, coordination and muscle condition of mice, a vertical 2 mm diameter smooth metal rod was fixed in the platform. Mouse grasped the wire with the two forepaws only, started the timer when the mouse was released. When the mouse fell off, the timer was stopped immediately and the hanging time was recorded. If the mouse could hang for more than 10 min, it was placed back in the cage. Falling mice were allowed to try up to two more times, at 2-minute intervals, and the maximum hanging time was recorded for further analysis *(Aartsma-Rus and van Putten,2014)*.

### Electrode implantation and video-electroencephalogram (v-EEG) recordings

Continuous v-EEG monitoring was performed for 1 week (7 days/week, 24 h/day).Screws (1*2 mm; Plexon, Dallas, TX, USA) were used as intracranial EEG electrodes and implanted 1 or 11 weeks post-TBI. One recording electrode was positioned anterior to the craniectomy window, and another was located contralaterally to the region. Reference and ground electrodes were placed in the ipsilateral and contralateral occipital bone, respectively (Fig. 2A). Mice were placed in Plexiglass cages and connected via tether to the v-EEG recording system (NT 9200 with a sample rate of 1000 Hz, Thermo Nicolet Corporation, USA, Fig. 2B). A modified “Seizure Severity Score” was used as a criterion for response to pentylenetetrazol (PTZ) *(Semple, et al.,2017; Van Erum, et al.,2019)*: 0 (no response), 1 (behavioral arrest), 2 (mouth and face twitches), 3 (head nodding and forelimb clonus), 4 (rearing), 5 (generalized convulsive seizure), 6 (continual generalized seizures), and 7 (status epilepticus resulting in death). The frequency of EEG abnormalities and severity of epileptic seizures were manually reviewed and scored based on both EEG and video recordings.

### Cresyl Violet Staining

Sections were put onto gelatin-covered adhesive glass slides, stained with cresyl violet for 5 min, and dehydrated in a series of 70%, 80%, 90%, and 100% alcohol for a few seconds each. Next, sections were cleaned twice with xylene for 2 min each. Images were acquired using an EVOS FL Auto 2 Imaging System (Thermo Fisher Scientific). Cresyl violet-stained coronal sections were imaged and histological lesion volumes were quantified with ImageJ software *(An, et al.,2016)* (http://imagej.nih.gov). The percentage of injured volume was calculated using the formula previously described by An *et al*. *(An, et al.,2016)*: [(VC−VL)/VC] × 100%, where VC is the volume of the contralateral hemisphere of the lesion, and VL is the volume of the ipsilateral hemisphere of the lesion.

### Immunofluorescence Staining

Brain slices and primary neurons were post-fixed with 4% PFA, washed three times with PBS, incubated for 30 min with 0.3% Triton, and then blocked with 1% goat serum for 1h to avoid binding of non-specific antibodies. Sections were then incubated at 4°C overnight with the following primary antibodies: anti-K_Na_1.1 (1:300; TA326566, Origene, Rockville, MD, USA), anti-NeuN, which is specific to nuclei and perinuclear cytoplasm of most of the neurons in the central nervous system of mammals and no glial cells (1:300; 26975-1-AP, Proteintech, Rosemont, IL, USA), anti-GABA-synthesizing enzyme glutamic acid decarboxylase 67 (GAD67) (1:300; ab213508, Abcam), anti-vesicular glutamate transporters1 (VGLUT1) (1:300; ab227805, Abcam). Secondary antibodies conjugated to Alexa Fluor 488 or 594 (1:1000; Invitrogen) were applied for 1.5h at room temperature. Slides were mounted with Fluoroshield containing DAPI (Ab104139; Abcam) and observed with a confocal microscope (LSM710; Zeiss, Oberkochen, Germany).

### Western Blotting

Fresh brain tissues were removed around the lesion and were lysed with radioimmunoprecipitation assay buffer (R0010, Solarbio) containing protease inhibitors and phosphatase inhibitors, and then centrifuged at 12,000 rpm at 4 °C for 10 min. Protein concentrations were determined using a Pierce™ BCA Protein Assay Kit (23227, Thermo Fisher Scientific). Immunoblotting was performed with the following antibodies: anti-GAD67 (1:500), anti-VGLUT1 (1:500), Anti-K_Na_1.1 (1:500), and tubulin (1:1000), β-actin (1:1000). Horseradish peroxidase-conjugated goat anti-mouse or goat anti-rabbit IgG (1:10000; Applygen) was used as the secondary antibody, and the signal was visualized using Super ECL Plus substrate (P1050, Applygen).

### Slice Electrophysiology

At 14 days post-injury (dpi 14), WT-Sham, WT-TBI, *kcnt1^−/−^*-Sham, and *kcnt1^−/−^*-TBI mice were deeply anesthetized with isoflurane and decapitated. Brain tissues were immediately submerged in ice-cold (0-4 °C) and oxygenated (95% O_2_ and 5% CO_2_) cutting solution containing (in mM): 126 NaCl, 25 NaHCO_3_, 10 D-glucose, 3.5 KCl, 1.5 NaH_2_PO_4_, 0.5 CaCl_2_, 10 MgCl_2_ (pH 7.35 and 340 mOsm). The brain was cut into 300 μm coronal slices with a Leica VT1000S. After incubating slices in fresh artificial cerebrospinal fluid (ACSF) containing (in mM): 126 NaCl, 3.5 KCl, 1.0 MgCl_2_, 2.0 CaCl_2_, 1.5 NaH_2_PO_4_, 25 NaHCO_3_, and 10 D-glucose; pH 7.3-7.4 and 310 mOsm) at 37 °C for 30 min, they were allowed to recover at room temperature for another 1 hour and then transferred to a submerged recording chamber located on an upright microscope (BX51WI; Olympus), which was perfused with oxygenated ACSF at a rate of 2-2.5 ml/min at the room temperature (22-23 °C). Whole-cell recordings were performed on cortical neurons visualized using infrared differential interference contrast microscopy using a Multiclamp 700B (Molecular Devices, San Jose, CA, USA) and Clampex 10.5 software (Molecular Devices). Data were acquired at 10 kHz and low-pass filtered at 1 kHz. Recording electrodes (resistance of 3-6 MΩ) were prepared from borosilicate glass capillaries (BF150-86-75; Sutter Instruments, Novato, CA, USA) using a vertical pipette puller (PC-100; Narishige, Tokyo, Japan). The pipette solution contained (in mM): 136 K-gluconate, 17.8 HEPES, 1 EGTA, 0.6 MgCl_2_, 4 ATP, 0.3 GTP, 12 creatine phosphate (pH 7.3 adjusted with KOH and osmolarity maintained at 290-300 mOsm).

As previously described *(Cheng, et al.,2022; Shore, et al.,2020)*, electrophysiological parameters recorded in our experiments included intrinsic membrane properties, and synaptic activities. We first divided neurons into excitatory pyramidal neurons (PNs) and non-fast spiking interneurons (INs) according to different morphological characteristics. PNs usually have clear apical and basal dendrites while INs are featured with small body size and obscure apical dendrite. Next, we utilized the electrophysiological parameters described in previous studies *(Avermann, et al.,2012; Shore, et al.,2020)* to distinguish PNs and INs. PNs showed obvious AP adaption with a wide AP half-width during AP trains. INs were confirmed if their maximum mean firing rates were in the range of 30-60 Hz, and with a larger input resistance and smaller AP half-width than PNs. Protocols were reported in a previous study *(Shore, et al.,2020)*. The threshold amplitude of sEPSCs was above 5 pA, while sIPSCs were two or three times that of baseline standard deviation (SD). The excitation/inhibition ratio = (sEPSC frequency * charge)/ (sEPSC frequency * charge + sIPSC frequency * charge).

### Cultured Cell Electrophysiology

Whole-cell recordings were performed on primary cortical neurons in parallel on the same day (day 13-16 in vitro), and viewed with an infrared differential interference contrast microscope with an Olympus 40× water immersion lens (BX51WI; Olympus, Tokyo, Japan) at room temperature (22-23°C). Standard extracellular solution contained the following (in mM): 140 NaCl, 2.4 KCl, 10 HEPES, 10 glucose, 4 MgCl_2_, and 2 CaCl_2_ (pH 7.35 and 320 mOsm). The internal solution, protocols, and determination of electrophysiological parameters were conducted as described above for slice recordings.

## Statistical Analysis

All results were presented as mean ± standard deviation (SD). Statistical analyses were done using GraphPad Prism 8 or SPSS 22 (IBM, Armonk, NY, USA). All data distributions were assessed with the Shapiro-Wilk test. Unpaired two-tailed Student’s t test, Kruskal-Wallis test, one-way ANOVA with Tukey’s post hoc tests, and two-way ANOVA with Sidak’s post hoc tests were used to determine statistical significance (p < 0.05).

## Acknowledgments

The authors thank Dr. Susan Campbell (University of Alabama at Birmingham, Department of Cell, Developmental and Integrative Biology) for helpful commentary on the manuscript and language editing.

## Author contributions

All authors contributed to the study conception and design. Material preparation and data collection were performed by Ru Liu and Lei Sun. The original draft was written by Ru Liu. Formal analysis, visualization, review and editing were performed by Du Le and Xi Guo. Manuscript revision and preparation of the response to reviewers were performed by Lulu Yu. Additional whole-cell patch-clamp experiments and data analysis were performed by Peng Pei. Data curation, review and editing, project administration, resources, and supervision of the research were performed by Qun Wang and Jianping Wu. All authors read and approved the final manuscript.

## Data Availability

The datasets generated during and/or analyzed during the current study are available from the corresponding author on reasonable request.

## Ethics approval

All procedures were approved by the Animal Care and Use Committee of Capital Medical University in accordance with the Guide for the Care and Use of Laboratory Animals. The studies involving human participants were reviewed and approved by the Medical Ethics Committee of Tiantan Hospital, Capital Medical University. Written informed consent to participate in this study was provided by the participants’ legal guardian/next of kin.

